# Androgen Promotes Differentiation of PLZF^+^ Spermatogonia pool via Indirect Regulatory Pattern

**DOI:** 10.1101/297846

**Authors:** Jingjing Wang, Jinmei Li, Yunzhao Gu, Qin Xia, Weixiang Song, Xiaoyu Zhang, Yang Yang, Wei Wang, Hua Li, Kang Zou

## Abstract

Androgen signaling plays a pivotal role in spermatogenesis, but the molecular mechanisms underlying androgen action in this process are unclear. Specifically, it is unknown if the androgen receptor (AR) is expressed in germ cells. Thus it’s interesting to reveal how androgen induces differentiation of spermatogonial progenitor cells (SPCs) in the niche. Here we observed the AR is primarily expressed in pre-spermatogonia of mice 2 days post partum (dpp), absent before spermatogenesis onset, and then expressed in surrounding Sertoli cells. Then we examined a regulatory role of the AR in spermatogenesis using a SPCs-Sertoli cells co-culture system, and demonstrated that androgen negatively regulated *Plzf* (the gene for stemness maintenance of SPCs). Additionally, we identified Gata2 as a target of AR in Sertoli cells, and demonstrated that Wilms tumor 1 (WT1) and β1-integrin as two putative intermediate molecules to transfer differentiation signals to SPCs, which was further verified using androgen pharmacological-deprivation mice model. These results demonstrate a regulatory pattern of androgen in SPCs niche in an indirect way via multiple steps of signal transduction.

## Introduction

Spermatogonial stem cells are the original source for developing functional sperms in male testis, which reside in the basement membrane and being under the control of their microenvironment. Androgens are produced by testicular Leydig cells and functions via binding and activation of its receptor AR in cytoplasm. Activated AR is subsequently imported into nucleus after release of heat shock protein and functions as a transcription factor ^1, 2^. Numerous target genes of AR have been identified, involving in many biological processes such as gonad development^3^ and tumorigenesis^4^. However, the exact biological mechanism of AR in spermatogenesis is not fully understood, and if AR is expressed in testicular germ cells is still unclear. Some earlier studies have detected AR expression in germ cells of mouse testis ^5, 6^. Other studies suggested that AR was presented only in the somatic cells in rodent testes ^7^. It has been shown that germ cell specific AR knockout mice still had normal sperm ^8^ but conditional deletion of AR in Leydig and Sertoli cells caused spermatogenesis defects ^9, 10^. These results suggests that AR expression in Sertoli cells, Leydig cells and perivascular myoid cells may participate in spermatogenesis via interacting with Sertoli cells surrounding spermatogonia^11^. However, Sycp1-driven Cre for AR deletion in germ cells was used in the study mentioned above^8^, which only indicates AR is not required in germ cells since meiosis onset. The exact mechanism between androgen and spermatogenesis still needs to be further investigated.

Promyelocytic Leukemia Zinc Finger (*Plzf*) is an important transcription suppressor gene for SSCs maintenance. It was first discovered by its association with acute promyelocytic leukemia ^12^, and was subsequently characterized as an undifferentiated marker for SSCs in rodents^13^, primates ^14^ and livestock ^15^. Loss of *Plzf* did not impact spermatogonia formation in embryonic and neonatal stages, but led to progressive and significant deficiency of SSCs after neonatal life and finally caused infertility ^13, 16^, indicating its critical role in SSCs maintenance. Moreover, PLZF expression was detected in spermatogonia A_s_, A_pr_ and A_al_, not restricted in SSCs ^17^. Thus, PLZF^+^ population represents undifferentiated spermatogonia pool in testis, and PLZF is an important factor for maintenance of this pool ^18, 19^. Although the interaction of androgen and PLZF has not been reported in germ cells, many evidence from prostate tumorigenesis studies suggest interaction between AR and PLZF. For example, *Plzf* has been shown to repress prostate tumorigenesis and its expression can be inhibited by androgen antagonist, bicalutamide ^20^. In prostate cancer cell line PCa cells, PLZF was identified as a repressor of AR as well as an activator of regulated in development and DNA damage responses 1 (REDD1), which suppressed mTORC1 ^21^. AR was characterized as a critical transcriptional factor in prostate tumorigenesis ^4^, and mTORC1 has been found to participate in EMT (Eepithelial mesenchymal transition) in prostate cancer ^22^. Thus, PLZF functions as tumor suppressor and interacts with AR in prostate cancer system, but it’s uncertain whether similar links exist in germ line. In rodent testis, Sertoli cells in base membrane form niches to protect SSCs and to regulate their fates ^23^, and many surface proteins, such as cadherins and integrins, have been identified as functional components in this microenvironment ^24^. Many of these molecules have been reported to be AR responsive and thus regulate the fate of SSCs ^25^, but the mechanism is still largely unknown.

To understand androgen’s role in Sertoli cells, it’s necessary to focus on the Wilms’ tumor 1 gene (*Wt1*), which is specifically expressed in mouse Sertoli cells and is required for Sertoli cell lineage maintenance ^26, 27^. Moreover, WT1 functions as a suppressor of *Ar* ^28^. Thus, we wonder whether WT1 participates in the regulation of spermatogenesis mediated by androgen signal.

To elaborate androgen’s regulatory mechanism in spermatogenesis, we studied AR expression pattern during postnatal testis development using a monoclonal antibody. Weak AR signal was detected in pre-spermatogonia of 2 dpp testes but not in surrounding somatic cells, however, from 3 dpp all signals were detected exclusively in somatic cells. Spermatogenesis starts from about 5 dpp ^29^, so the possibility that germ cells need AR for spermatogenesis is eliminated. In vitro experiments further confirmed this conclusion, thus we investigated the indirect regulation pattern of androgen on SSCs differentiation via Sertoli cells using a SSCs-Sertoli cells co-culture system. Two transcription factors Gata2 and WT1 were identified as AR’s downstream targets in Sertoli cells. Moreover, a surface protein β1-integrin was subsequently regulated by WT1 and may function as an intermediate molecule to transfer androgen signal from Sertoli cells to SSCs to regulate spermatogenesis. In all, this study demonstrates the complicated regulation pattern of androgen in spermatogenesis in SSCs niche, which requires the cooperation of Sertoli cells and many intermediate molecules to regulate *Plzf* expression in undifferentiated spermatogonia population.

## Results

### PLZF^+^ spermatogonia pool keeps steady during testis development

Due to PLZF is a specific marker for undifferentiated spermatogonia, population size of PLZF^+^ cells in mouse testis determines the capacity of self-renewal and spermatogenesis in seminiferous tubules. Our immunohistochemistry results demonstrated that from neonatal to adult testes, all PLZF^+^ cells exclusively located at basement membrane with strong nucleus staining [Fig.1A-D], or a few oblate pairs of cells in adult testes, which are A_pr_ spermatogonia [Fig.1D inset]. These PLZF^+^ cells were surrounded by Sertoli cells, which formed the niches for SSCs. Strong PLZF intensity in 5 dpp testis indicated high expression level of PLZF at this stage. In testes from 5 to 42 dpp, the percentage of PLZF^+^ cells in seminiferous tubules was declined [Fig.1Q] due to continuous production of differentiated germ cells. However, the total number of PLZF^+^ cells was approximately consistent [Fig.1R], indicating a stable pool of undifferentiated spermatogonia during testicular development.

**Figure 1.**
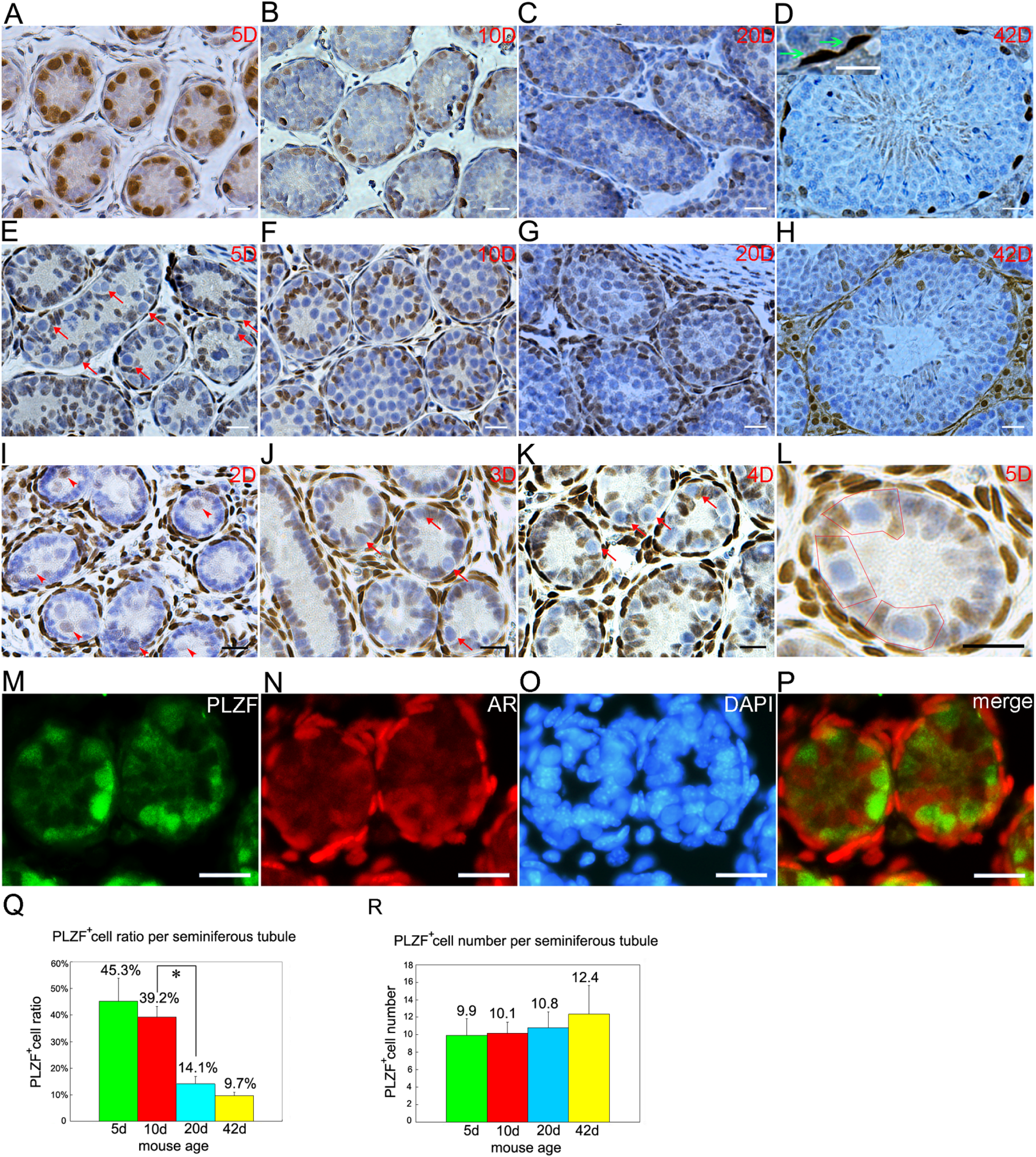
Expression patterns of PLZF and AR during testis development. PLZF staining was exclusively detected in spermatogonia localized in basal membranes of 5 dpp (A), 10 dpp (B), 20 dpp (C) and 42 dpp (D) testes (inset in D, representive of PLZF^+^ A_pr_ spermatogonia). AR staining in 5 dpp (E), 10 dpp (F), 20 dpp (G) and 42 dpp (H) testes was specifically detected in Sertoli cells in the first or second layers of seminiferous tubules. Weak AR staining was observed in pre-spermatogonia (red arrowheads) in the testicular lumen of 2 dpp testis, but not in Sertoli cells (I). In 3 dpp testis (J), pre-spermatogonia migrated to basement membranes and formed unique structures with surrounding Sertoli cells termed niches (red arrows in J, and identical structures were also found in 4 dpp and 5 dpp testes, red arrows in E and K), and AR staining was detected in Sertoli cells but not in germ cells. K. AR staining in 4 dpp old testis, L. a representive of niche in 3 dpp testis, in which SSCs were embraced by AR^+^ Sertoli cells (red frame). AR and PLZF dual immunofluorescence exhibited AR signals and PLZF signals were not overlapped in 4 dpp testis (M. PLZF, N. AR, O. DAPI, P. merge). From neonatal to adult stages, ratio of PLZF^+^ cells in seminiferous tubules decreased (Q) but number of PLZF^+^ cells kept steady (R). Data represent means ± SD (*p < 0.01). Bar=20μM.

### AR expression in testis indicates an indirect regulation pattern in promoting spermatogenesis

To investigate the function of androgen during spermatogenesis, we examined AR expression pattern in neonatal (5 dpp), juvenile (10 dpp), puberty (20 dpp) and adult (42 dpp) testes using immunohistochemistry. Intensive AR signals were detected in Sertoli cells, Leydig cells and myoid cells but not in germ cells [Fig.1 E-H], consistent with previous report that postnatal testicular germ cells lacked endogenous AR ^7^. In 5 dpp testes, moderate AR staining was detected in Sertoli cells localized at basement membrane [Fig.1 E]. Both AR^+^ Sertoli cells number and AR signal intensity were obviously increased in 10 dpp testes, and some AR^+^ Sertoli cells moved to the second layers [Fig.1 F], and then formed AR^+^ layers in 20 dpp testes [Fig.1 G]. In 42 dpp testis, those AR^+^ second layers disappeared and AR^+^ cells distributed in different layers of seminiferous tubules [Fig.1 H]. In rodent Sertoli cells, AR expression was reported to start from 3-5 dpp ^30, 31^, thereupon we examined AR expression in 2, 3 and 4 dpp testes, and detected AR signal in Sertoli cells only from 3 dpp testis and signal intensity grew stronger from 5 dpp [Fig.1I-K]. We also noticed Sertoli cells did not express AR in 2 dpp testis as literature reported ^31^, but we unexpectedly observed weak AR signal in pre-spermatogonia localized in the cord of testes [Fig.1I]. Interestingly, AR signal was absent in the pre-spermatogonia of 3 dpp testes, at which time point some of these pre-spermatogonia had already migrated from cord to basal membrane [Fig. 1J]. Meanwhile, Sertoli cells began to express AR and two AR^+^ Sertoli cells embraced one pre-spermatogonium to form a special structure nominated as ‘niche’ [Fig.1J-L], implying AR in Sertoli cells commenced to function in microenvironment. It’s known that spermatogenesis starts from 3-5 dpp and differentiated spermatogonia can be observed at 6 dpp ^29^. Dual immunofluorescent staining also exhibited no overlap of AR and PLZF signals in 4 days testes [Fig.1 M-P]. Collectively, AR expression pattern in Sertoli cells demonstrates their close connection with PLZF^+^ cells, but PLZF^+^ population is unable to be directly stimulated by androgen since they lack of endogenous AR when spermatogenesis starts.

### Blocking of androgen by antagonist inhibits SSCs differentiation

To investigate the role of androgen in spermatogenesis, a co-culture system of testicular cells was established. Testes from 5 dpp mice (this time spermatogenesis just begins) were harvested and digested to single cells (composed of SSCs, differentiating spermatogonia and testicular somatic cells) and cultured on dishes. Somatic cells attached to gelatin and formed flat layers within 12 hours’ culture, and undifferentiated spermatogonia population including SSCs attached on these layers [Fig.2A]. This total testicular cells co-culture system can be maintained in vitro for 4 passages (16 days) at least [Fig.2B]. Moreover, this co-culture system stably expressed germ line markers and somatic markers [Fig.S1A-V], and was sensitive to dihydrotestosterone (DHT) or bicalutamide (an efficient AR antagonist) stimulation [Fig.S1W], indicating spermatogonia (including SSCs) and Sertoli cells were well maintained in this system and can be used to mimic spermatogenesis in vitro. To study androgen’s function in spermatogenesis in vitro, total testicular cells from 5dpp testes were plated on dishes, and 16 hours later DHT was supplied to culture medium, and 2 hours later bicalutamide was supplied to corresponding samples. After 48-hour incubation, more A_al_ stage spermatogonia were observed in DHT treated group while more SSCs clusters remained in bicalutamide treated group [Fig.2C-E]. RT-PCR revealed DHT stimulation caused decreased expression levels of *Id4*, *Gfra1* and *Plzf* [Fig. 2F], indicating reduced size of undifferentiated spermatogonia populations including A_s_, A_pr_ and A_al_, while expression level of c-kit was significantly increased [Fig. 2F]. Notably, expression of *Tex14*, a marker of intercellular bridge existing in both mitotic and meiotic divisions, namely from A_pr_, A_al_ spermatogonia, to spermtocytes ^32^, was up-regulated after DHT treatment [Fig. 2F]. Combined with observations above, we concluded that DHT caused a reduced size of undifferentiated spermatogonia populations, and accordingly proposed that androgen promoted differentiation of spermatogonia in this in vitro system. Results from western blot revealed expression levels of AR displayed opposite responses to those of PLZF after DHT/bicalutamide treatment [Fig. 2G]. Additionally, we observed that DHT aggravated down-regulation of PLZF expression induced by siRNA [Fig. 2H], which further confirmed that DHT promoted SSCs differentiation. On the contrary, Thy1-MACS purified SSCs cultured on gelatin or mitotically inactivated STO feeder layer (negative of AR expression) showed no obvious phenotype change after either DHT or bicalutamide treatment (data is not shown), suggesting Sertoli cells were necessary for SSCs differentiation during androgen stimulation.

**Figure 2.**
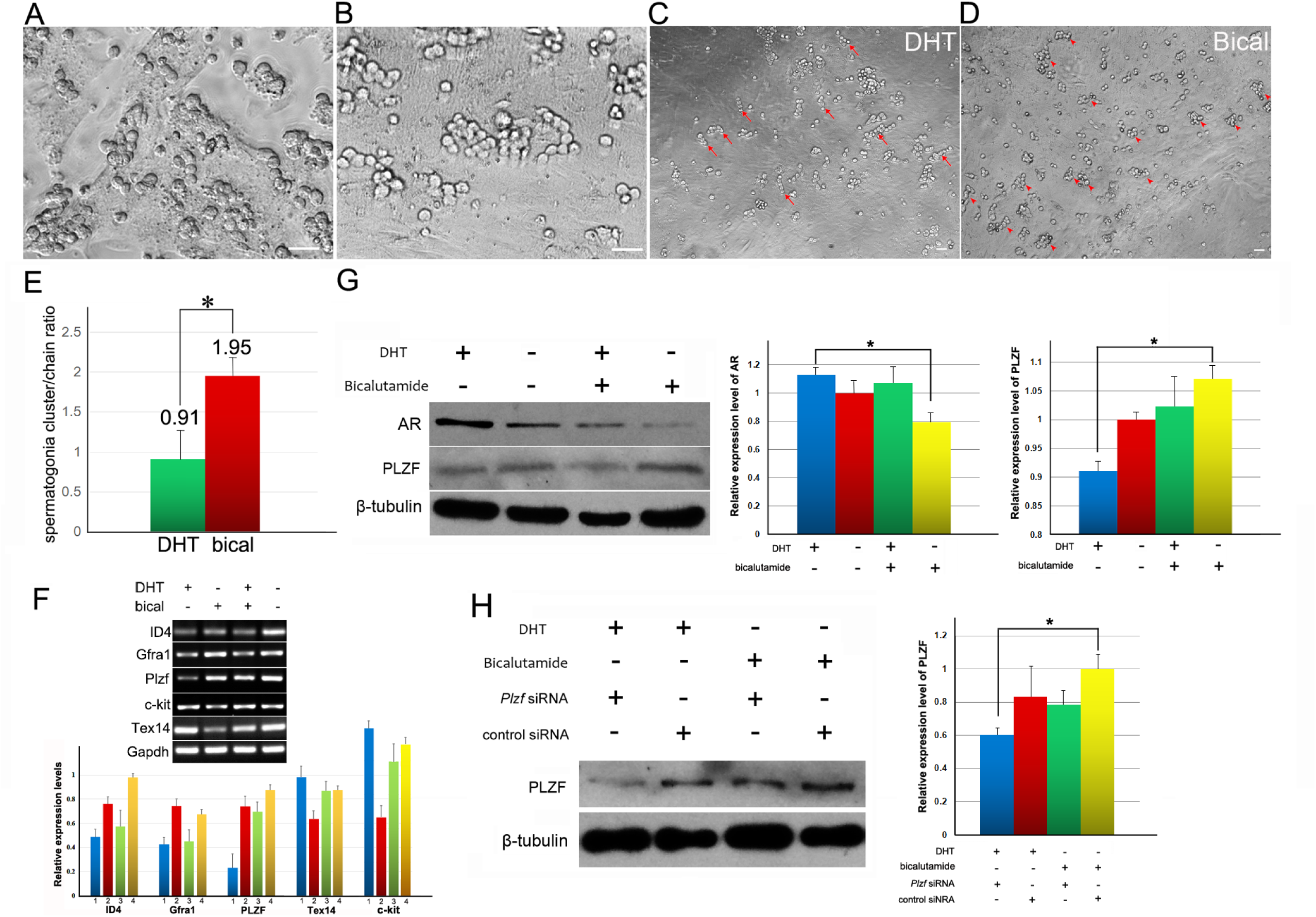
Impact of androgen on SSCs differentiation in SSCs-testicular somatic cells co-culture system. Total testicular cells isolated from testes of 5-day mice were maintained in vitro, and SSCs grew well on the layers formed by somatic cells (A). The morphology of co-culture system after two passages was exhibited (B). More A_al_ spermatogonia (red arrows) were observed in DHT treat group (C) while more clusters (red arrow heads) formed in bicalutamide treat group (D). The ratio of spermatogonia cluster/chain was calculated after DHT or bicalutamide treatment in co-culture system (E). Expression levels of *Id4*, *Gfra1*, *Plzf*, *c-kit*, *Tex14* in SSCs-Sertoli cells co-culture system after DHT and/or bicalutamide treatment were evaluated by RT-PCR (F). Impact on expression of AR and PLZF in co-culture system caused by DHT and bicalutamide treatment was determined by western blot (G). DHT aggravated the decline of PLZF expression caused by PLZF siRNA, while bicalutamide alleviated the knockdown effect (H).

### AR indirectly regulates *Wt1* in Sertoli cells

To reveal regulatory mechanism of AR on spermatogenesis, we used ChIP-seq to screen the target genes of AR in purified Sertoli cells from 5 dpp mouse testis. In the candidates of targets, we noticed that *Gata2* promoter was a binding region of AR (p-value <0.01 by MACS ^33^) [Fig.3A]. This result provided an important cue, since Gata2 has been identified as a factor to promote WT1 expression by binding the enhancer of *Wt1* ^34^, and WT1 is a pivotal transcription factor for spermatogenesis in Sertoli cells ^35^. Then we did ChIP-qPCR to validate the binding of AR on *Gata2* [Fig. 3 B], and observed that AR negatively regulated expression of Gata2 and WT1 after DHT stimulation or AR overexpression in Sertoli cells [Fig.3C]. Consistently, knockdown of Gata2 caused down-regulated WT1 expression [Fig.3D]. Taken together, these results implied that AR indirectly regulated WT1 expression via Gata2.

**Figure 3.**
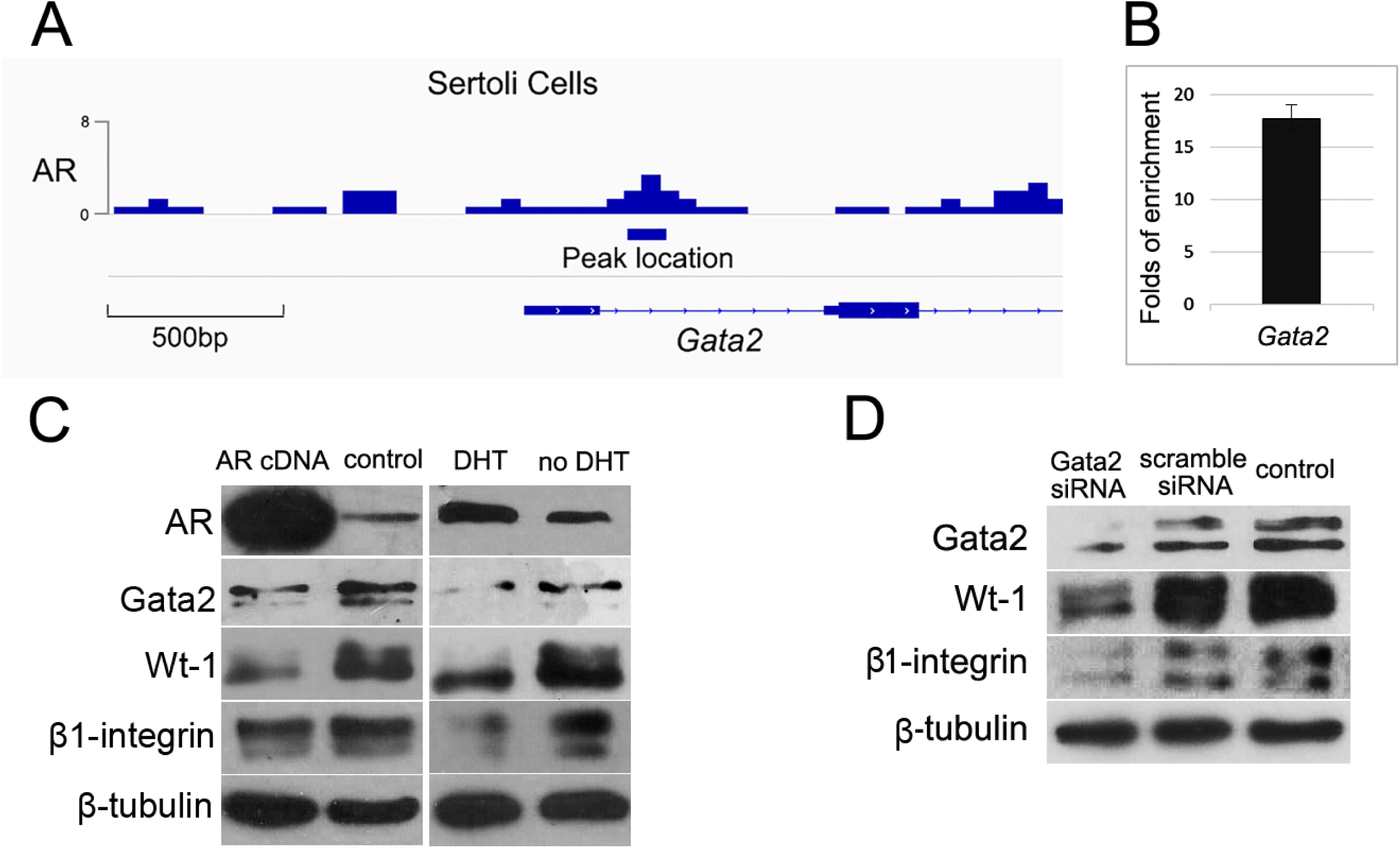
Screening for target genes of AR in Sertoli cells. The representative view of AR binding sites (visualized by Integrative Genomics Viewer [http://software.broadinstitute.org/software/igv/]) at *Gata2* promoter region (A). ChIP-qPCR validated the binding of AR in the promoter region of *Gata2*: fold enrichment by antibodies against AR relative to control IgG was presented as mean value ± SD of two replicates (B). Overexpression of AR or DHT treatment caused down-regulated expression levels of Gata2, WT1 and β1-integrin in Sertoli cells (C). Knockdown of Gata2 using siRNA led to decreased expressions of WT1 and β1-integrin (D).

### The *β1-integrin* gene is a direct target of WT1 in Sertoli cells

As a pivotal transcription factor for spermatogenesis, WT1 involves multiple functions in Sertoli cells. Thus, we used ChIP-seq again to screen the potential target genes of WT1 in Sertoli cells. Notably, β1-integrin gene was identified as a direct binding target of WT1 [Fig. 4A]. Besides as a subunit of laminin receptor, β1-integrin is essential for SSCs homing since deletion of β1-integrin in SSCs or in Sertoli cells impaired SSCs homing ^24^, indicating its signal function in SSCs niche. After validated the binding of WT1 on β1-integrin gene by ChIP-qPCR [Fig. 4B], we further examined whether WT1 regulates β1-integrin expression at protein level. Knockdown of WT1 expression in testis somatic cells (SSCs removed) caused down-regulated β1-integrin expression [Fig. 4C], and WT1 overexpression strengthened β1-integrin’s signal [Fig. 4D], suggesting β1-integrin expression was regulated by WT1. Moreover, we observed that androgen signal negatively regulated expression of β1-integrin in Sertoli cells [Fig. 4E], and disturbance of Gata2 expression displayed the same impact on β1-integrin expression [Fig. 3C], which were consistent with aforementioned results that AR negatively regulated Gata2 by binding on its promoter region, and consequently interfered in WT1 expression. Then we interfered in β1-integrin expression in co-culture system using siRNA and observed decreased expression level of PLZF [Fig.4F], implying the indispensable role of β1-integrin in SSCs maintenance. Collectively, we preliminarily found a cue from androgen signal to surface protein β1-integrin, via intermediate factor Gata2 and WT1 [Fig. 4G].

**Figure 4.**
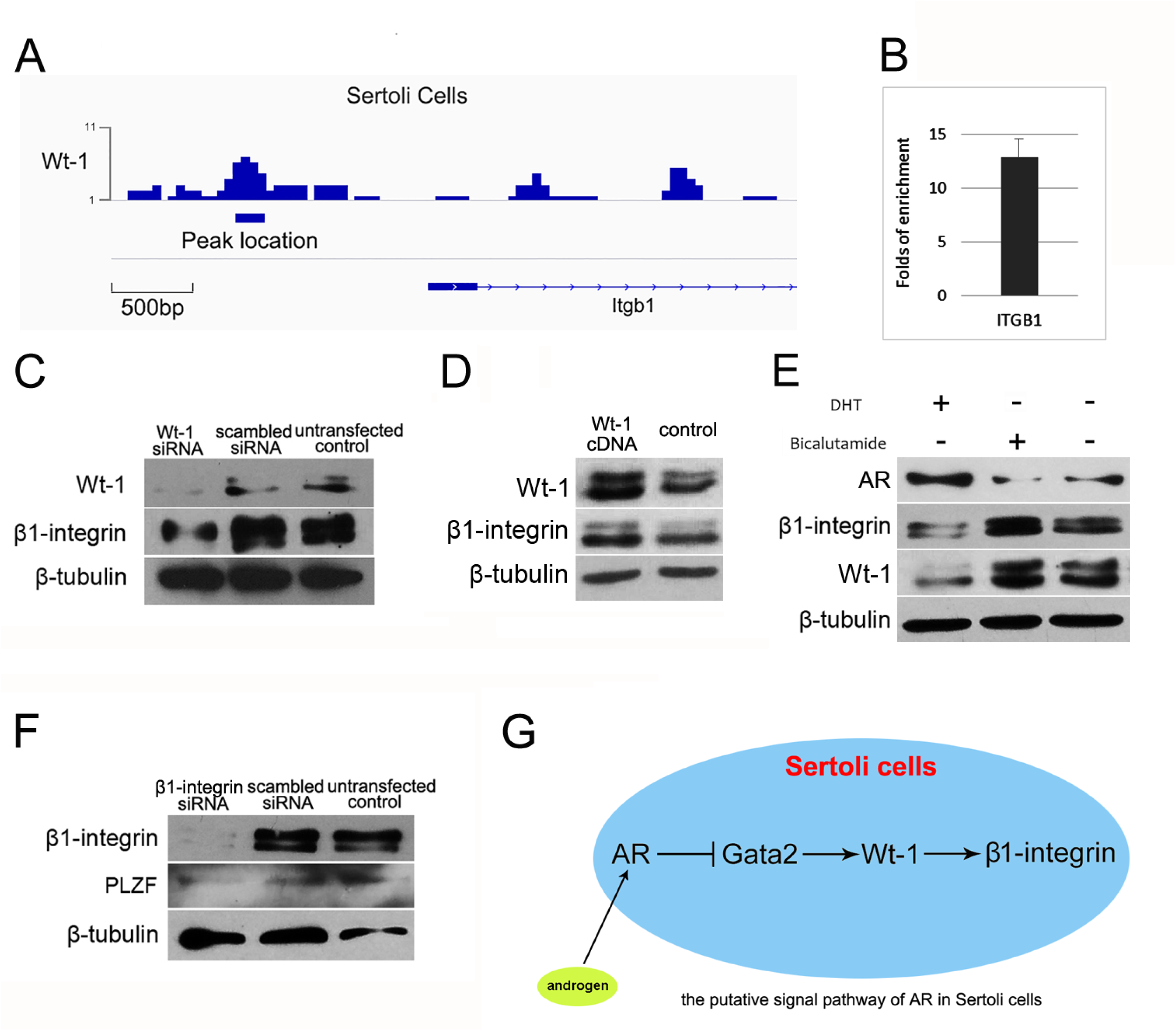
WT1 is an intermediate molecule to transmit androgen signal to downstream target β1-integrin. The representative view of WT1 binding sites (visualized by Integrative Genomics Viewer [http://software.broadinstitute.org/software/igv/]) at *β1-integrin* promoter region (A). ChIP-qPCR validates the binding of WT1 in the promoter region of *β1-integrin*: fold enrichment by antibodies against WT1 relative to control IgG was presented as mean value ± SD of two replicates (B). Knockdown of WT1 in testicular somatic cells decreased β1-integrin expression (C), and transfection of *Wt1* cDNA enhanced β1-integrin expression (D). Androgen caused down-regulation of WT1 and β1-integrin in co-culture system (E). Knockdown of β1-integrin in this co-culture system led to decreased expression level of PLZF (F). An illustration hypothesizes the potential signal pathway from AR to β1-integrin in Sertoli cells (G).

### β1-integrin on Sertoli cells may function as an intermediate molecule to regulate SSCs differentiation

The important role of β1-integrin in vivo has been proven in Sertoli cells specific β1-integrin conditional knockout mice ^24^. Here we used in vitro co-culture system to preliminarily investigate the role of β1-integrin on Sertoli cells for SSCs maintenance. Because primary Sertoli cells have limited passage capacity in vitro, and knockout of β1-integrin in Sertoli cells caused serve damage, we infected primary Sertoli cells with β1-integrin shRNA or control shRNA lentivirus for 72 hours to attenuate β1-integrin expression and used them as feeder layers for Thy1^+^ SSCs (Knockdown efficiency of β1-integrin shRNA was evaluated in Fig. S2J-L). After co-culture for 48 hours, number of SSCs on Sertoli cells infected with β1-integrin shRNA lentivirus was significantly less than that on control shRNA infected Sertoli cells [Fig. 5A], and western blot confirmed SSCs on β1-integrin interfered feeder expressed reduced levels of SSCs markers [Fig. 5B], indicating that β1-integrin on Sertoli cells affects maintenance of SSCs in this co-culture system.. Then we hypothesize that β1-integrin on Sertoli cells must interact with some molecules on SSCs to regulate their fates. To screen the molecules interacted with β1-integrin, expression of integrins and cadherins in SSCs were examined b**y** RT-PCR [Fig. 5C]. We focused on E-cadherin as a candidate since SSCs highly and specifically express E-cadherin. Subsequently, the binding of β1-integrin and E-cadherin was validated in adult mouse testis lysates using immunoprecipitation. E-cadherin was pulled down by the antibody against-β1-integrin, and vice versa [Fig. 5D], implying that E-cadherin on SSCs binds with β1-integrin in physiological condition. However, different from E-cadherin as a specific undifferentiated spermatogonia marker, β1-integrin is expressed on both spermatogonia and Sertoli cells ^36, 37^, which is consistent with our results from IF [Fig.5E,G and Fig. S3]. Thus, it’s not clear that E-cadherin on SSCs binds with β1-integrin on Sertoli cells or on SSCs themselves. Therefore, thy1^+^ SSCs were used for IP [Fig. 5H], and the result eliminates the possibility that E-cadherin and ITGB1 on SSCs binds on themselves. These results suggest that β1-integrin on Sertoli cell is essential for SSCs maintenance, and probably binds with E-cadherin on SSCs as a putative interaction partner.

**Figure 5.**
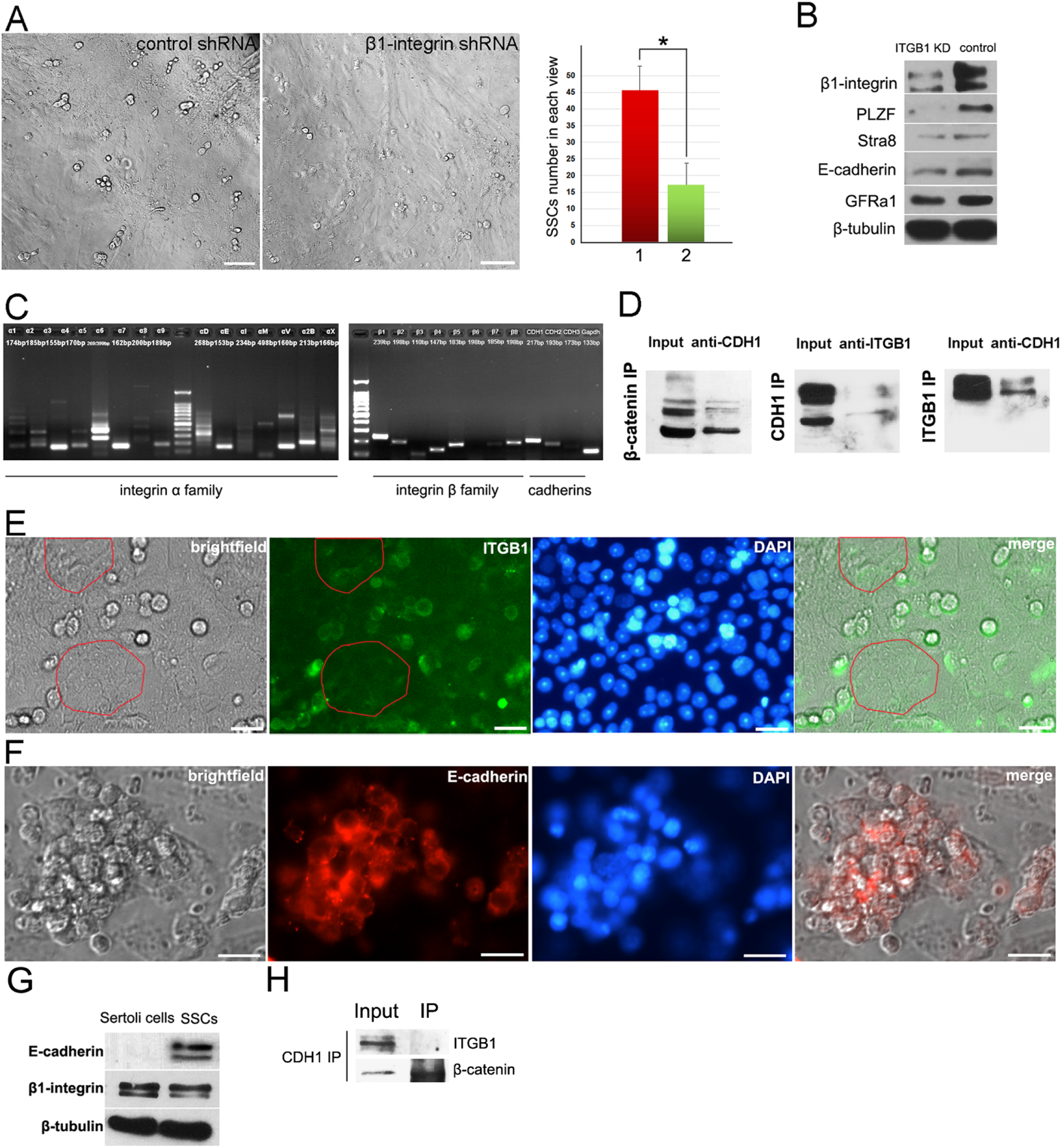
Identification of β1-integrin as a putative interactive molecule between SSCs and Sertoli cells. Thy1^+^ SSCs were co-cultured with Sertoli cells infected with control shRNA or β1-integrin shRNA for 48 hours (A), and the expression levels of germline markers of SSCs were detected (B). RT-PCR revealed expression profiles of integrins and cadherins in SSCs at mRNA level (C). The binding of E-cadherin and β1-integrin was detected in adult mouse testis cell lysates using immunoprecipitation, of which the binding of β-catenin and E-cadherin was used as positive control (D). Expression of β1-integrin (E) and E-cadherin (F) in SSCs and co-cultured Sertoli cells was determined by IF. Purified Sertoli cells or Thy-1^+^ SSCs were lysed to detect the expression of β1-integrin and E-cadherin using western blot (G). Thy-1^+^ SSCs were lysed for co-IP to detect the endogenous interaction of β1-integrin and E-cadherin (H).

### Pharmacologically androgen deprived mice are lack of differentiated germ cells but enriched with hyper-proliferated PLZF^+^ and β1-integrin^+^ spermatogonia

To further study the role of androgen on spermatogenesis, 33 6-week old male mice were intraperitoneally injected with bicalutamide, since mice in this age had mature Sertoli cells, stable AR expression and continuous spermatogenesis. Bicalutamide recipient mice exhibited retarded body weight growth and smaller testis size compared to control group [Fig. 6A-B]. Histological results revealed the severe deficiency of differentiated germ cells (spermatocytes, spermatids and sperms) in seminiferous tubules [Fig. 6E-F]. Moreover, we observed reduced number of AR^+^ cells [Fig. 6D] and decreased expression level of AR [Fig.6G, H and R], but obvious increased number of PLZF^+^ cells [Fig.6C] and expression level of PLZF [Fig.6I, J and R] in bicalutamide recipient testes, indicating that deprivation of androgen did not impair undifferentiated spermatogonia population, but resulted in blockage of germ cell differentiation and expansion of undifferentiated spermatogonia population. Notably, in many seminiferous tubules of bicalutamide recipients, accumulated spermatogonia formed a second layer [Fig.6K], which were identified as PLZF negative [Fig.6M], β1-integrin and c-kit positive [Fig.6N-O], and mitotically active [Fig.6P-Q] spermatogonia. These results indicated androgen deprivation blocked spermatogenesis at the step that ckit^+^β1-integrin^+^ spermatogonia differentiate to more differentiated populations, and consequently led to accumulation of undifferentiated spermatogonia, including PLZF^+^ population [Fig. 7A]. Western blot revealed enhanced expressions of Gata2, WT1 in bicalutamide recipients [Fig. 6R], which was consistent with results from in vitro system that androgen regulated Sertoli cells’ function in spermatogenesis via Gata2 and WT1. Enhanced expressions of PLZF and β1-integrin resulted from accumulated spermatogonia, and decreased expressions of Sycp3, Claudin-11 and Nectin-2 were consequence of lost meiotic germ cells caused by androgen deprivation [Fig.6R]. These results implied that blockage of androgen in adult mouse testis inhibited the differentiation from spermatogonia to spermatocytes and promoted SSCs self-renewal involving multiple signal transductions in the niche. We will focus on how β1-integrin of Sertoli cells interacts with SSCs to regulate SSCs differentiation in the future work.

**Figure 6.**
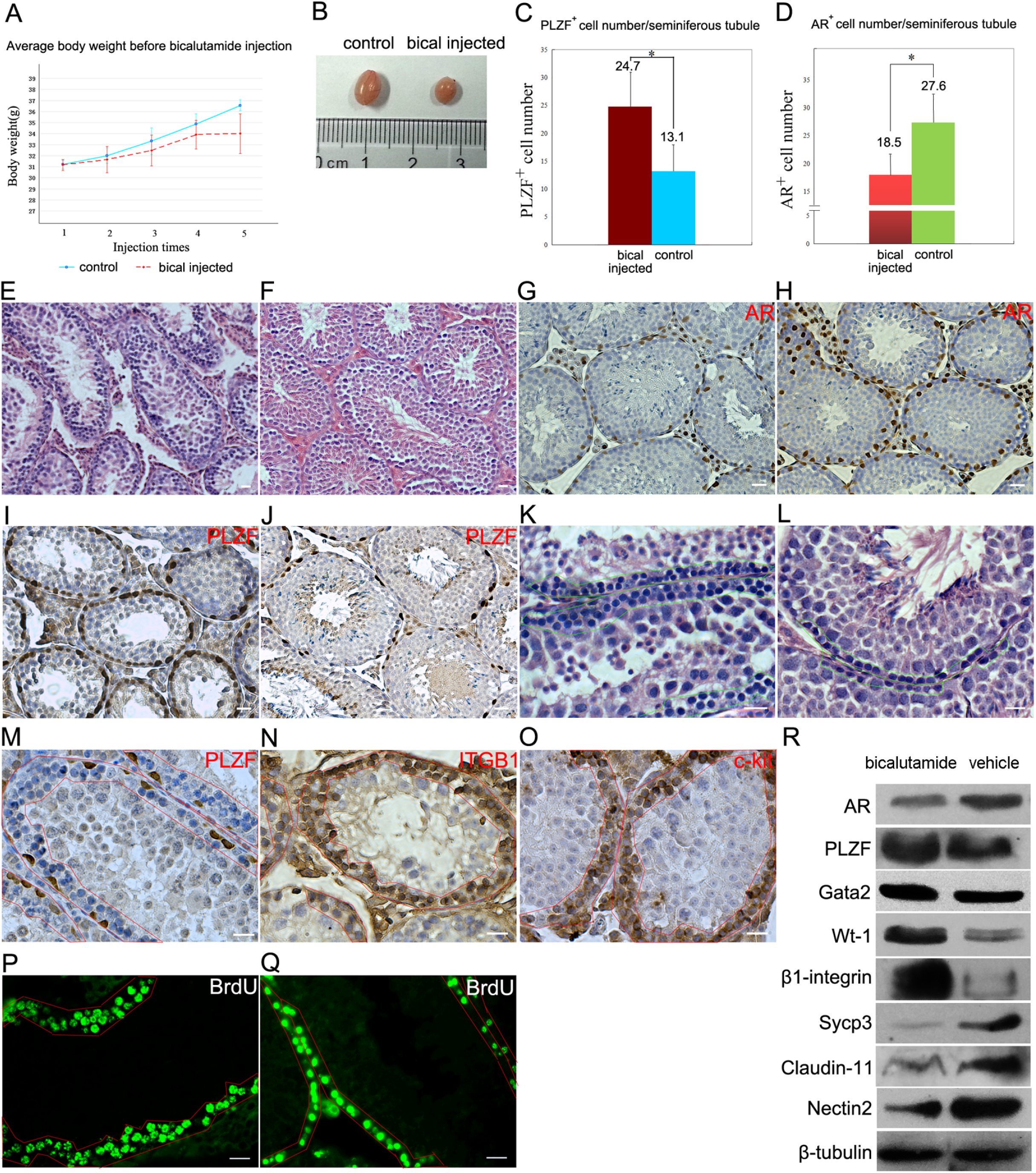
Bicalutamide recipient mice lack of spermatocytes and spermatids, but possess hyper-proliferative SSCs. Bicalutamide recipient mice (n=33) exhibited retarded body weight growth (A) and smaller testis size (B). Histology demonstrated significantly reduced spermatocytes and spermatids in seminiferous tubules of bicalutamide recipients (E) compared with those of control (F). IHC staining displayed reduced number of AR^+^ cells (G) and increased number of PLZF^+^ cells (I) in bicalutamide recipients than in controls (H and J). A statistical result indicated a augmented population of PLZF^+^ cells (C) and a remarkable decline of AR^+^ cell population (D) in seminiferous tubules of bicalutamide recipients, data represent means ± SD (*p < 0.01). Accumulated spermatogonia formed two layers in bicalutamide recipient seminiferous tubules (green frame in K), while wild type seminiferous tubule possessed only one layer of spermatogonia (green frame in L). Most of the spermatogonia in those two layers were β1-integrin^+^ (N), c-kit^+^ (O) PLZF^−^ (M). BrdU assay indicated they were mitosis active (P), as those in wild type (Q). Spermatogonia layers were enclosed by red frames in M, N, O, P and Q. Bicalutamide recipient testes displayed down-regulated expression of *Ar*, *Sycp3*, *Claudin-11*, *WT1*, *Nectin-2*, and up-regulated expression of *Plzf*, *Gata2*, *Wt1* and *β1-integrin* (R).

**Figure 7.**
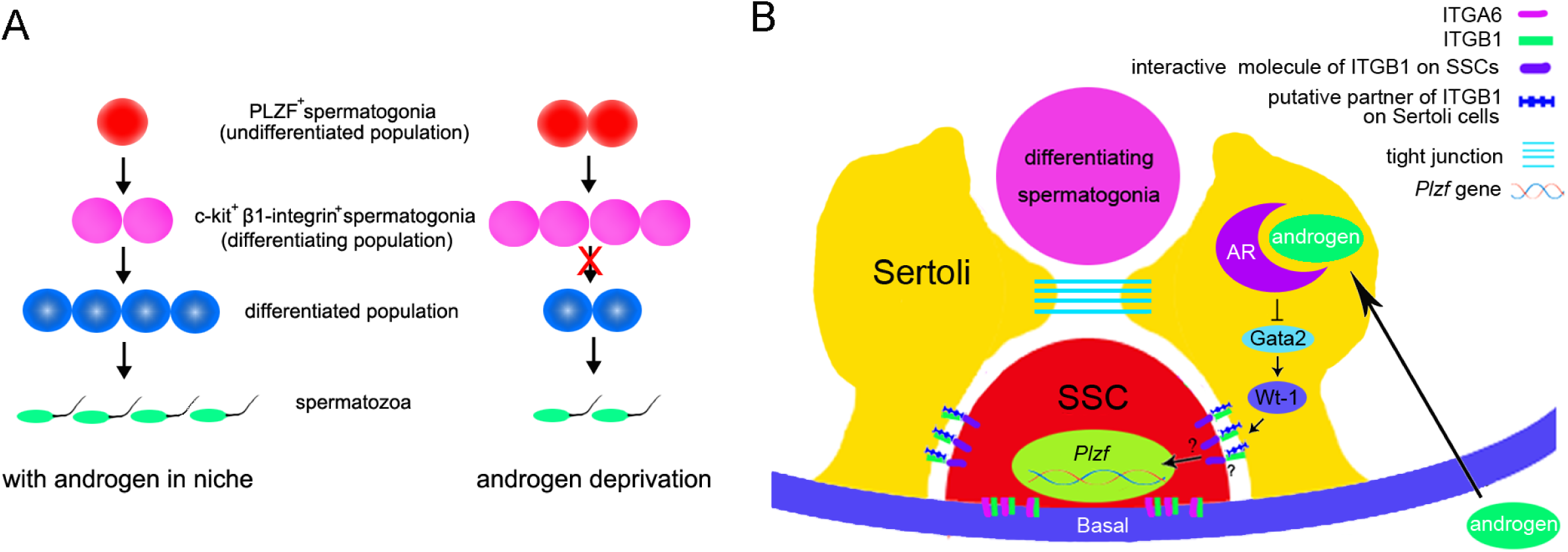
The hypothesis of putative regulation mechanism of spermatogenesis induced by androgen. Androgen deprivation may block spermatogenesis at the step that β1-integrin^+^c-kit^+^ population transits to more differentiated germ cells, causing accumulation of undifferentiated spermatogonia (A). Androgen produced by Leydig cells interacts with AR in Sertoli cells to regulate down-stream target genes, and subsequently transfer differentiation signals to SSCs, and finally turns off the differentiation inhibitor gene *Plzf* via multiple steps. *Gata2* was identified as a target gene of AR, and *Wt1* was activated by Gata2 to regulate downstream gene *β1-integrin* in Sertoli cells. How β1-integrin on Sertoli cells influences SSCs’ fates by regulation of *Plzf* expression is unknown (B).

## Discussion

Our results exhibit SSCs lack of endogenous AR after 3 dpp, which is consistent with the dispensable role of AR in differentiated germ cells from AR germ cell conditional knockout mice. Interestingly, some studies detected AR expression in germ cells of fetal testis ^38^, and we examined weak AR signal in pre-spermatogonia of 2 dpp old testes. Therefore, we prefer the opinion that SSCs do not need endogenous AR after neonatal stage, and the function of AR in embryonic and new born stages is still need to be revealed.

Testis Zinc Finger Protein (TZFP), a homolog of PLZF, was reported to co-repress the activated AR ^39^ and may participate in hormone signal pathway to regulate spermatogenesis ^40^. However, IHC staining showed PLZF and AR staining were distributed in distinct cell types after neonatal stage, indicating no direct interaction between these two transcription factors during spermatogenesis. Based on our data, we hypothesize that androgen signal transduction in spermatogenesis spatially can be summarized into three steps [Fig. 7B]: the first step is AR activation in Sertoli cells induced by androgen, including androgen-AR interaction and activation of AR’s downstream genes to produce signal molecules; second step is the communication between Sertoli cells and SSCs, involving some surface or transmembrane proteins; the last one is signal transduction to SSCs nucleus to turn off PLZF and start differentiation. Our results suggest that AR inhibits gata2 expression in Sertoli cells by binding on its promoter region, leading to subsequent decrease of WT1 expression, and WT1 binds and regulates β1-integrin in Sertoli cells, which interacts with SSCs as a putative signal molecule to regulate SSCs differentiation. WT1 is able to bind the promoter of *Ar* to inhibit AR expression ^28^. We revealed AR’s inhibitory role on WT1 expression via interacting with Gata2, indicating a feedback regulation on WT1 expression. Moreover, we proposed β1-integrin on Sertoli cells as a regulatory molecule on SSCs differentiation. Routinely, β1-integrin combines with another subunit, i.e. α1-integrin, α6-integrin, to form a complex. On SSCs surface β1-integrin binds α6-integrin to form laminin receptor, which is essential for SSCs maintenance. Nevertheless, it’s not clear that α6-integrin binds with β1-integrin in Sertoli cells^41^, thus we are interesting in searching for the interactive molecule of β1-integrin on Sertoli cells in future study. Moreover, although our data demonstrated that E-cadherin on SSCs binds β1-integrin on Sertoli cells, it’s not clear E-cadherin really functions as a signal molecule to regulate SSCs fates in responding to β1-integrin on Sertoli cells, especially loss of E-cadherin in SSCs has been proven with unimpaired homing capacity. However, we can not conclude E-cadherin is redundant in SSCs, since SSCs enrich other types of cadherins which may compensate the role of E-cadherin, and that study also noticed that E-cadherin mutation in SSCs led slightly reduced colony formation after transplantation ^24^. Moreover, E-cadherin was reported to be functional in transformation of SSCs into pluripotent status ^42^. Thus, in future work we will focus on the link of E-cadherin and SSCs fate, and will also screen β1-integrin’s other binding partners on SSCs surface related to signal reception and transmission to SSCs nucleus.

Two important junction proteins Clauding-11 and Nectin-2, which were reported to be significantly impacted in testes of AR knockout mouse ^25^, were down-regulated in bicalutamide recipient testes. Claudin-11 is a pivotal component of tight junction (TJ) barrier and closely related to spermatogenesis, and its deficiency caused infertility ^43^. *Ar* deletion in Sertoli cells resulted in down-regulation of Claudin-3 (a homolog of Claudin-11 and a component of TJ junction) and a subsequent increase of BTB (blood-testis barrier) permeability ^44^. Further study revealed that Claudin-11 replaced Claudin-3 during spermatocyte translocation in spermatogenesis ^45^. Therefore, we inferred that androgen deprivation caused the blockage of spermatogonia differentiation and a consequent accumulation of SSCs. Reduced Claudin-11 expression after androgen deprivation indicated loose of TJ junction. Whether Claudin-11 interacts with β1-integrin is also an interesting question. Down-regulation of Nectin-2 after androgen deprivation from our experiments exhibited contradicting result to that from AR knockout mice ^25^. As a pivotal anchor junction protein, Nectin-2 expression was mainly detected in Sertoli cells and elongated spermatids, and *Nectin-2* knockout mice had normal sperm titers but exhibited infertility phenotype due to serve spermatozoa malformation ^46^, suggesting the major role of Nectin-2 is for sperm mature. Thus, the dramatic decline of Nectin-2 expression in our androgen deprivation model was caused by blockage of spermatogonia differentiation into spermatocytes and spermatids.

This study reveals androgen signals pathway in Sertoli cells which promoting differentiation of PLZF^+^ spermatogonia population. Gata2, WT1 and β1-integrin were identified as pivotal intermediate molecules in this process. In the future, we will focus on the communication pattern between Sertoli cells and SSCs, in particular β1-integrin’s role in regulation of SSCs fates.

## Materials and Methods

### Animals

CD-1 mice were supplied by Comparative Medicine Centre of Yangzhou University and all the procedures for animal experiments were approved by the ethical committee at Nanjing Agricultural University.

### Isolation and culture of total testicular cells or SSCs

Testicular cells were extracted from testes of 5 dpp mice generally following previous protocol ^47^ with minor modifications. Testes were cut into small particles and followed by collagenase IV and trypsin digestion. Undigested collagen was removed using micropippetor after centrifugation, and cell pellet was resuspended in culture medium and then transferred to gelatin coated plates for incubation. For SSCs sorting, resuspended cells after two-enzyme digestion were filtered with 70-µm cell filter and subsequently incubated with mouse Thy-1.2 antibody coated magnetic beads (BD, Cat.551518). Thy1+ fraction was collected and cultured on mitotically inactivated STO feeder layer at 37 °C with 5% CO2. Culture medium for total testicular cells or SSCs was composed of 90% MEMa, 10% FBS (Gibco), 1 ng/ml bFGF (Sino Biological Inc., 10014-HNAE), 1 ng/ml EGF (Sino Biological Inc., 50482-MNAY), 1 ng/ml GDNF (Sino Biological Inc., 10561-HNCH), 10 ng/ml LIF (santa, sc-4378), 20 μg/ml transferrin (sigma, T8158), and 5 μg/ml insulin (aladdin, I113907). For androgen stimulation or deprivation, dihydrotestosterone (sigma, cat. 31573) or bicalutamide (sigma, PHR-1678) was added to the culture medium accordingly 48 hours before cell harvesting. To make the feeder layer, STO cells were incubated with 10 µg/ml mitomycin C (Roche, cat. 101074090001) at 37 °C with 5% CO2 for 2 hours and plated on gelatin coated plates.

### Sertoli cell isolation

A modified method was used to isolate Sertoli cells from testes of 6 dpp mice^48^. Briefly, testes were encapsulated and seminiferous tubules were pooled and digested with 0.5mg/mL collagenase Ⅳ. After centrifugation, the supernatant containing the Leydig cells was pipetted out. The tubules were further digested with 1 mg/mL collagenase Ⅳ and 10 μg/mL DNase to detach peritubular cells. Digestion was monitored under microscopy. The supernatant containing peritubular cells after centrifugation was discarded. Finally, single cell suspension was obtained after trypsin digestion and blocked by FBS. The cell pellet was resuspended in DMEM/F12 media supplemented with 2.5% FBS, 5% horse serum, 2mM L-Gln, 100U/ml penicillin, 1uM sodium pyruvate, and then plated. After 48h hypotonic treatment was performed to remove germ cells as described^49^, and the purity of Sertoli cell was assessed by morphology and immunofluorescent staining of WT1(purity of Sertoli cells>95%) [Fig.S2A-D,I].

### RNA extraction and RT-PCR

Total RNA was isolated using TRIzol (TIANGEN,DP405) and first-strand cDNA was synthesized using a PrimeScript RT Master Mix(Takara,RR036A)for the reverse transcription (RT)-PCR. The information of primers was listed in Table S1.

### ChIP-seq and ChIP-qPCR

To characterize genome-wide binding patterns of androgen receptor and WT1 in Sertoli cells, ChIP and input DNA libraries were performed as previously described^50^. Briefly, cells were cross-linked with 1% formaldehyde for 10 min at room temperature and formaldehyde was then inactivated by the addition of 125 mM glycine for 5min. Sonicated DNA fragments with 100–300 bp were pre-cleared and then immunoprecipitated with Protein A+G Magnetic beads coupled with Anti-Androgen Receptor antibody (06–680, Millipore). After reverse crosslinking, immunoprecipitated DNAs and input DNAs were end-repaired and ligated adapters to the DNA fragments using NEBNext Ultra End-Repair/dA-Tailing Module (E7442, NEB) and NEBNext Ultra Ligation Module (E7445, NEB). High-throughput sequencing of the ChIP fragments was performed using Illumina NextSeq 500 following the manufacturer’s protocols. The raw sequencing data were processed with trimmomatic (version 0.36) to filter low-quality reads ^51^. The resulting data were mapped using bowtie2 (version 2.2.9) to the UCSC mm10 genome reference ^52^. Peak detection was performed using the MACS peak finding algorithm (Model-based Analysis of ChIP-Seq; version 1.4.2) with parameters --nomodel --shiftsize 25 and the p-value cutoff set to 0.05 ^33^.

Primers for ChIP-qPCR:

GATA2-F: GCTAGAGAGTGCATTGGGGA

GATA2-R:AGTTCCTGGGGCTGCGAG

β1-integrin F:

β1-integrin R:

Control-F:TTAGGTGGCCTCAGATCCTC

Contro-R:CCTGCCTCTCTTTTGGACAG

### Bicalutamide injection

Bicalutamide injection was performed as described ^53^ with minor modifications: 6-week old CD1 male mice were intraperitoneally injected with bicalutamide (20 mg/kg) or DMSO (vehicle) once every other day for 4 times.

### Immunohistochemistry and immunofluorescence

Mouse testes were harvested and fixed in 4% neutral paraformaldehyde overnight, and subsequently dehydrated and embedded in paraffin. Histological sections were dewaxed and rehydrated in ethanol series, followed by microwave antigen retrieval in 0.01M citrate (pH=6.0) and methanol/H_2_O_2_ treatment at room temperature. After blocking with 5% goat serum, the slides were incubated with primary and biotin labeled secondary antibodies, respectively. Streptividin-HRP (Jackson Lab, 1:500) and DAB kit (Vector, sk4100) were used for visualization. Cells immunofluorescence was carried out as described ^54^. See antibodies information at Table S2.

### BrdU assay

Mice were intraperitoneally injected with BrdU (sigma, B5002, 50 mg/kg) 4 hours before testis harvesting. Testes were fixed in 4% neutral PFA and embedded with paraffin. For BrdU detection, sections were incubated with 2M HCl at 37 °C for 1 hour, and washed with 0.1M boric acid (pH= 8.5) for 3 times. The immunohistochemistry was performed as aforementioned (BrdU primary antibody, santa cruz, sc-32323, 1:200 dilution).

### Gene silence and overexpression

The validated siRNA or shRNA sequences (table 1) were synthesized by Shanghai Genepharma company. AR expression plasmid was a gift from Prof. Zijie Sun (School of Medicine, Stanford University). *Wt1* cDNA expression plasmid (EX-Mm24822-02) and *Itgb1* cDNA plasmid (MM1013-202859073) were purchased from GeneCopoeia and Dharmacon, respectively. Plasmids or siRNA oligos were transfected using lipofectamine 3000 (Lifetechnology).

**Table 1.**
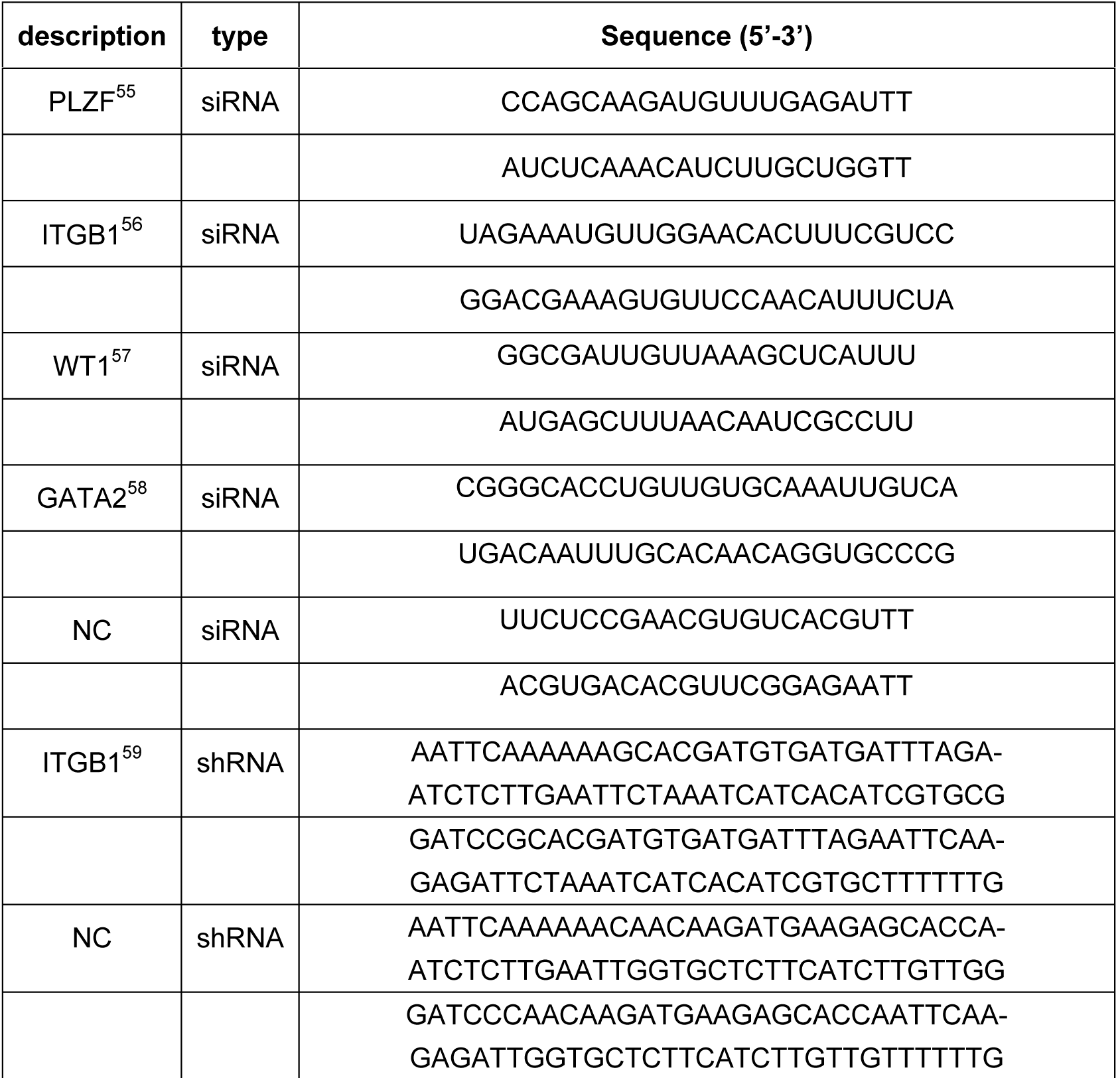
Sequence of small interfering or hairpin RNA

### Western blot and Co-IP

For western blotting, tissue and cell lysates were separated in SDS PAGE gel and transferred to nitrocellulose membrane, which was subsequently blocked with 5% skim milk at 4 °C overnight and then incubated with primary antibodies at room temperature for 1 h. Then the membranes were incubated with HRP-conjugated goat anti rabbit or mouse IgG (santa cruz, 1:3000) and ECL, and finally exposed the film. See antibodies information at Table S2. For Co-IP, lysates of adult mice testis or SSCs of 6 dpp mice were prepared using a standard protocol (Beyotime,P0013), and 0.5mg total proteins diluted by TBST were incubated with mouse anti-itgb1 antibody and rabbit anti-cdh1 antibody respectively and rocked at 4°C for overnight. The pre-washed protein (A+G) sepharose beads (Beytime,P2012) were added afterwards and incubated at 4°C for 4 hours. The beads were then washed three times with TBST, pelleted and boiled in 1XSDS loading buffer, and finally analyzed by western blot. See antibodies information at Table S2.

### Statistical analysis

For cell counting, sections or immunofluorescent visual fields were selected randomly. Data were analyzed by Excel and was presented as mean ± SD (standard deviation). Statistical significance was determined by *t*-test.

## Acknowledgments

We thank Prof. Zijie Sun and Xiaodong Zhao’s suggestions for AR signal study and ChIP-seq data analysis. This work was supported by the National Natural Science Foundation of China (81200472/H0425) and Fundamental Research Funds for the Central Universities in China (KYTZ201602).

---

**Figure S1. SSCs-somatic cells co-culture system was able to be maintained in vitro for short term.** SSCs-somatic cells co-culture system stably expressed MVH (A. bright filed, B. MVH, C. dapi, D. merge), PLZF (E. bright filed, F. PLZF, G. dapi, H. merge), WT1 (I. bright filed, J. WT1, K. dapi, L. merge), AR (M. bright filed, N. AR, O. dapi, P. merge) and CD9 (Q. bright filed, R. WT1, S. dapi, T. merge). RT-PCR results exhibit the co-culture system expresses somatic markers (U. 1.*Gapdh*, 2.*Sox9*, 3. *Sma*, 4.*Coopu-tf*, 5. *Wt1*, 6.*Mvh*), and *Ar* (V.1) *Ar*, and germline markers (V. 2.*Scp3*, 3.*ckit*, 4.*integrin-aV*, 5.*Cd9*, 6.*integrin-1*, 7.*Plzf*, 8. *Mvh*) after 3 passages. Western blot result confirmed this co-culture system was sensitive to DHT or bicalutamide stimulation (W).

**Figure S2. Purified Sertoli cells were used for ChIP-seq analysis and co-culture.** Bright field (A), WT1 staining (B), DAPI (C) and merge (D) of purified Sertoli cells. WT1^+^ ratio of purified Sertoli cells was higher than 95% (I). Purified Sertoli cells were examined for β1-integrin expression by IF (E-H), and then were infected with β1-integrin shRNA lentivirus to interfere in endogenous expression of β1-integrin before co-culture with SSCs. Morphology of Sertoli cells 96 hours post infection (J), and infection efficiency was examined by expression of reporter gene RFP (red fluorescence protein) (K). Western blot revealed relative expression level of β1-integrin was approximately decreased by 80% after β1-integrin shRNA interfering (L).

Author contributions
Jinmei Li: Collection and/or assembly of data, data analysis and interpretation
Jingjing Wang: Collection and/or assembly of data, data analysis and interpretation
Yunzhao Gu: Collection and/or assembly of data, data analysis
Weixiang Song: Collection and/or assembly of data, data analysis
Xiaoyu Zhang: Collection and/or assembly of data
Yang Yang: Collection and/or assembly of data
Wei Wang: Data analysis and interpretation
Hua Li: Data analysis and interpretation
Kang Zou: Conception and design, financial support, manuscript writing, final approval of manuscript

## Reference

1 Balk SP. Androgen receptor as a target in androgen-independent prostate cancer. Urology 2002; 60:132–138.

2 Rommerts FF, Melsert R, Teerds KJ, De Rooij DG. Multiple regulation of Leydig cell functions. Growth Factors in Fertility Regulation 1991.

3 Krutskikh A, de Gendt K, Sharp V, Verhoeven G, Poutanen M, Huhtaniemi I. Targeted Inactivation of the Androgen Receptor Gene in Murine Proximal Epididymis Causes Epithelial Hypotrophy and Obstructive Azoospermia. Endocrinology 2011; 152:689–696.

4 Zhu C, Luong R, Zhuo M et al. Conditional Expression of the Androgen Receptor Induces Oncogenic Transformation of the Mouse Prostate. Journal of Biological Chemistry 2011; 286:33478–33488.

5 Kimura N, Mizokami A, Oonuma T, Sasano H, Nagura H. Immunocytochemical localization of androgen receptor with polyclonal antibody in paraffin-embedded human tissues. Journal of Histochemistry & Cytochemistry 1993; 41:671–678.

6 Vornberger W, Prins G, Musto NA, Suarez-Quian CA. Androgen receptor distribution in rat testis: new implications for androgen regulation of spermatogenesis. Endocrinology 1994; 134:2307–2316.

7 Takeda H, Chodak G, Mutchnik S, Nakamoto T, Chang C. Immunohistochemical localization of androgen receptors with mono- and polyclonal antibodies to androgen receptor. Journal of Endocrinology 1990; 126:17-NP.

8 Tsai M-Y, Yeh S-D, Wang R-S et al. Differential effects of spermatogenesis and fertility in mice lacking androgen receptor in individual testis cells. Proceedings of the National Academy of Sciences of the United States of America 2006; 103:18975–18980.

9 Xu Q, Lin H-Y, Yeh S-D et al. Infertility with defective spermatogenesis and steroidogenesis in male mice lacking androgen receptor in Leydig cells. Endocr 2007; 32:96–106.

10 Karel DG, Swinnen JV, Saunders PTK et al. A Sertoli Cell-Selective Knockout of the Androgen Receptor Causes Spermatogenic Arrest in Meiosis. Proceedings of the National Academy of Sciences of the United States of America 2004; 101:1327–1332.

11 Zhou Q, Nie R, Prins GS, Saunders PTK, Katzenellenbogen BS, Hess RA. Localization of Androgen and Estrogen Receptors in Adult Male Mouse Reproductive Tract. Journal of Andrology 2002; 23:870–881.

12 Wang ZY, Chen Z, Huang W et al. Problems existing in differentiation therapy of acute promyelocytic leukemia (APL) with all-trans retinoic acid (ATRA). Blood Cells 1993; 19:633–641; discussion 642-637.

13 Costoya JA, Hobbs RM, Maria B et al. Essential role of PLZF in maintenance of spermatogonial stem cells. Nature Genetics 2004; 36:551–553.

14 Hermann BP, Sukhwani M, Lin CC et al. Characterization, Cryopreservation, and Ablation of Spermatogonial Stem Cells in Adult Rhesus Macaques. Stem Cells 2007; 25:2330–2338.

15 Pramod RK, Mitra A. In vitro culture and characterization of spermatogonial stem cells on Sertoli cell feeder layer in goat (Capra hircus). J Assist Reprod Genet 2014; 31:993–1001.

16 Filipponi D, Hobbs RM, Ottolenghi S et al. Repression of kit Expression by Plzf in Germ Cells. Molecular and Cellular Biology 2007; 27:6770–6781.

17 Payne C, Braun RE. Histone lysine trimethylation exhibits a distinct perinuclear distribution in Plzf-expressing spermatogonia. Developmental Biology 2006; 293:461–472.

18 Lovelace DL, Gao Z, Mutoji K, Song YC, Ruan J, Hermann BP. The regulatory repertoire of PLZF and SALL4 in undifferentiated spermatogonia. Development 2016; 143:1893–1906.

19 Hobbs RM, Seandel M, Falciatori I, Rafii S, Pandolfi PP. Plzf Regulates Germline Progenitor Self-Renewal by Opposing mTORC1. Cell 2010; 142:468–479.

20 Feng J, Zhou W. Identification and characterization of PLZF as a prostatic androgen-responsive gene. Prostate 2004; 59:426–435.

21 Yang J, Su Q, Martina T et al. Molecular circuit involving KLK4 integrates androgen and mTOR signaling in prostate cancer. Proceedings of the National Academy of Sciences of the United States of America 2013; 110: E2572–E2581.

22 Chen XG, Cheng HY, Pan TF et al. mTOR regulate EMT through RhoA and Rac1 pathway in prostate cancer. Molecular Carcinogenesis 2015; 54:1086–1095.

23 Kanatsu-Shinohara M, Miki H, Inoue K et al. Germline niche transplantation restores fertility in infertile mice. Human Reproduction 2005; 20:2376–2382.

24 Kanatsu-Shinohara M, Takehashi M, Takashima S et al. Homing of Mouse Spermatogonial Stem Cells to Germline Niche Depends on β1Integrin. Cell Stem Cell 2008; 3:533–542.

25 Ruey-Sheng W, Shuyuan Y, Lu-Min C et al. Androgen receptor in sertoli cell is essential for germ cell nursery and junctional complex formation in mouse testes. Endocrinology 2006; 147:5624–5633.

26 Rio-Tsonis KD, Covarrubias L, Kent J, Hastie ND, Tsonis PA. Regulation of the Wilms’ tumor gene during spermatogenesis. Developmental Dynamics An Official Publication of the American Association of Anatomists 1996; 207: 372.

27 Myers M, Ebling FJ, Nwagwu M et al. Atypical development of Sertoli cells and impairment of spermatogenesis in the hypogonadal (hpg) mouse. Journal of Anatomy 2006; 207:797–811.

28 Zaia A, Fraizer G, Piantanelli L, Saunders G. Transcriptional regulation of the androgen signaling pathway by the Wilms’ tumor suppressor gene WT1. Anticancer Research 2001; 21:1–10.

29 McLean DJ, Friel PJ, Johnston DS, Griswold MD. Characterization of Spermatogonial Stem Cell Maturation and Differentiation in Neonatal Mice. Biology of Reproduction 2003; 69:2085–2091.

30 You L, Sar M. Androgen receptor expression in the testes and epididymides of prenatal and postnatal sprague-dawley rats. Endocr 1998; 9:253–261.

31 Bremner WJ, Millar MR, Sharpe RM, Saunders PT. Immunohistochemical localization of androgen receptors in the rat testis: evidence for stage-dependent expression and regulation by androgens. Endocrinology 1994; 135:1227–1234.

32 Greenbaum MP, Yan W, Wu MH et al. TEX14 is essential for intercellular bridges and fertility in male mice. Proceedings of the National Academy of Sciences of the United States of America 2006; 103:4982–4987.

33 Zhang Y, Liu T, Meyer CA et al. Model-based analysis of ChIP-Seq (MACS). Genome Biology 2008; 9:: R137.

34 Furuhata A, Murakami M, Ito H et al. GATA-1 and GATA-2 binding to 3’ enhancer of WT1 gene is essential for its transcription in acute leukemia and solid tumor cell lines. Leukemia 2009; 23:1270–1277.

35 Wang XN, Li ZS, Ren Y et al. The Wilms Tumor Gene, Wt1, Is Critical for Mouse Spermatogenesis via Regulation of Sertoli Cell Polarity and Is Associated with Non-Obstructive Azoospermia in Humans. PLoS Genetics 2013; 9:e1003645.

36 Salanova M, Stefanini M, Curtis ID, Palombi F. Integrin receptor alpha 6 beta 1 is localized at specific sites of cell-to-cell contact in rat seminiferous epithelium. Biology of Reproduction 1995; 52:79–87.

37 Salanova M, Ricci G, Boitani C, Stefanini M, De GS, Palombi F. Junctional contacts between Sertoli cells in normal and aspermatogenic rat seminiferous epithelium contain alpha6beta1 integrins, and their formation is controlled by follicle-stimulating hormone. Biology of Reproduction 1998; 58:371–378.

38 Merlet J, Racine C, Moreau E, Moreno SG, Habert R. Male fetal germ cells are targets for androgens that physiologically inhibit their proliferation. Proceedings of the National Academy of Sciences of the United States of America 2007; 104:3615–3620.

39 Kaufmann S, Sauter M, Schmitt M et al. Human endogenous retrovirus protein Rec interacts with the testicular zinc-finger protein and androgen receptor. Journal of General Virology 2010; 91:1494–1502.

40 Furu K, Klungland A. Tzfp Represses the Androgen Receptor in Mouse Testis. PLoS ONE 2013; 8: e62314.

41 Cheng CY, Lie PP, Mok KW et al. Interactions of laminin β3 fragment with β1-integrin receptor: A revisit of the apical ectoplasmic specialization-blood-testis-barrier-hemidesmosome functional axis in the testis. Spermatogenesis 2011; 1:174–185.

42 An J, Zheng Y, Dann CT. Mesenchymal to Epithelial Transition Mediated by CDH1 Promotes Spontaneous Reprogramming of Male Germline Stem Cells to Pluripotency. Stem Cell Reports 2016; 8:446–459.

43 Mazaud-Guittot S,., Meugnier E,., Pesenti S,. et al. Claudin 11 deficiency in mice results in loss of the Sertoli cell epithelial phenotype in the testis. Biology of Reproduction 2010; 82:202–213.

44 Meng J, Holdcraft RW, Shima JE, Griswold MD, Braun RE. Androgens regulate the permeability of the blood–testis barrier. Proceedings of the National Academy of Sciences of the United States of America 2005; 102:16696–16700.

45 Smith BE, Braun RE. Germ Cell Migration Across Sertoli Cell Tight Junctions. Science 2012; 338:798–802.

46 Mueller S, Rosenquist TA, Takai Y, Bronson RA, Wimmer E. Loss of Nectin-2 at Sertoli-Spermatid Junctions Leads to Male Infertility and Correlates with Severe Spermatozoan Head and Midpiece Malformation, Impaired Binding to the Zona Pellucida, and Oocyte Penetration. Biology of Reproduction 2003; 69:1330–1340.

47 Wu J, Zhang Y, Tian GG et al. Short-type PB-cadherin promotes self-renewal of spermatogonial stem cells via multiple signaling pathways. Cellular Signalling 2008; 20:1052–1060.

48 Ks VDW, Johnson EW, Dirami G, Dym TM, Hofmann MC. Immunomagnetic isolation and long-term culture of mouse type A spermatogonia. Journal of Andrology 2001; 22:696–704.

49 Galdieri M, Ziparo E, Palombi F, Russo MA, Stefanini M. Pure Sertoli Cell Cultures: A New Model for the Study of Somatic—Germ Cell Interactions. Journal of Andrology 1981; 2:249–254.

50 Zhang XL, Wu J, Jian W et al. Integrative epigenomic analysis reveals unique epigenetic signatures involved in unipotency of mouse female germline stem cells. Genome Biology 2016; 17: 162.

51 Bolger AM, Lohse M, Usadel B. Trimmomatic: a flexible trimmer for Illumina sequence data. Bioinformatics 2014; 30:2114–2120.

52 Li H. Aligning sequence reads, clone sequences and assembly contigs with BWA-MEM. 2013; 1303.

53 Jin R, Yamashita H, Yu X et al. Inhibition of NF-kappa B signaling restores responsiveness of castrate resistant prostate cancer cells to anti-androgen treatment by decreasing androgen receptor variants expression. Oncogene 2015; 34:3700–3710.

54 Zou K, Yuan Z, Yang Z et al. Production of offspring from a germline stem cell line derived from neonatal ovaries. Nature cell biology 2009; 11:631–636.

55 Schefe JH, Menk M, Reinemund J et al. A novel signal transduction cascade involving direct physical interaction of the renin/prorenin receptor with the transcription factor promyelocytic zinc finger protein. Circulation Research 2006; 99: 1355.

56 Zeng Q, Guo Y, Liu Y et al. Integrin-β1, not integrin-β5, mediates osteoblastic differentiation and ECM formation promoted by mechanical tensile strain. Biological Research 2015; 48:1–8.

57 Jing Z, Wei-Jie Y, Yi-Feng ZG. Down-regulation of Wt1 activates Wnt/β-catenin signaling through modulating endocytic route of LRP6 in podocyte dysfunction in vitro. Cellular Signalling 2015; 27:1772–1780.

58 Schang AL, Granger A, Quérat B, Bleux C, Cohen-Tannoudji J, Laverrière JN. GATA2-induced silencing and LIM-homeodomain protein-induced activation are mediated by a bi-functional response element in the rat GnRH receptor gene. Molecular Endocrinology 2013; 27:74–91.

59 Mori H, Lo AT, Inman JL et al. Transmembrane/cytoplasmic, rather than catalytic, domains of Mmp14 signal to MAPK activation and mammary branching morphogenesis via binding to integrin β1. Development (Cambridge, England) 2013; 140:343–352.

